# Monitoring for fisheries or for fish? Declines in monitoring of salmon spawners continue despite a conservation crisis

**DOI:** 10.1101/2024.12.01.626233

**Authors:** Emma M Atkinson, Bruno S Carturan, Clare P Atkinson, Andrew W Bateman, Katrina Connors, Eric Hertz, Stephanie J Peacock

## Abstract

Monitoring of salmon in Pacific Canada has been declining for decades. Counts of spawning salmon are critically important, enabling researchers to quantify impacts of stressors, identify when and where management interventions are required to avoid extirpations, and evaluate the efficacy of recovery efforts. These data are more important now than ever, as uniquely adapted spawning populations underlie salmon resilience and their ability to adapt to climate change, and fine-scale data can inform sustainable fishing opportunities including the revitalization of terminal fisheries. We revisit the state of Pacific salmon spawner data from the Yukon to southern British Columbia. Almost two-thirds of salmon populations that were historically monitored have no reported estimates in 2014-2023 - the worst decade for data since broadscale spawner surveys began in the 1950s. We found a positive association between the number of populations monitored and landed value for three of the five Pacific salmon species, suggesting that monitoring is, to some extent, motivated by the information needs of commercial fisheries management. We recommend aligning monitoring objectives and strategic investments to improve monitoring outcomes for salmon, ecosystems, and the communities that depend on them. We emphasize data stewardship, as ensuring access to these baseline data is a cornerstone for rebuilding wild Pacific salmon.

## Introduction

Collecting baseline, fishery-independent data is a pillar of sustainable fisheries management, providing the foundation for evidence-based decisions about the conservation and management of aquatic species (Hilborn and Walters 1992; Punt et al. 2020; Punt 2023). While commercial fisheries data are important components of sustainable fisheries management frameworks, relying solely on fishery-dependent data is broadly acknowledged as insufficient to meet sustainable fisheries objectives (Walsh et al. 2018; Free et al. 2020; Ovando et al. 2021). In a recent example underlining this point, Canada’s Pacific salmon fishery was forced to withdraw from the Marine Stewardship Council certification program in 2022, in large part due to a lack of sufficient data to assess salmon abundance trends (Raincoast Conservation Foundation 2019; Marine Stewardship Council 2022). The history of colonial fisheries and their management in Canada is punctuated with consequential failures resulting from insufficiently considering the biology of fished species and the harvest levels they can sustain. From the 1990s collapse of Atlantic cod (Hutchings and Rangeley 2011) to the mid-2000s decline of Fraser sockeye in the Pacific (Peterman and Dorner 2011; Castaneda et al. 2020), retrospective reviews emphasized the importance of decision-making grounded in robust, data-driven science. Maintaining up-to-date and quality-controlled datasets, required for rigorous science, is crucial to effectively managing fish populations. In the absence or decline of commercial fisheries, as in the case of wild salmon in Pacific Canada (Walters et al. 2019), baseline data remain instrumental to stewarding populations through stressors such as climate change and habitat degradation. Multiple types of data inform efforts to study, monitor, and manage Pacific salmon. The federal department of Fisheries and Oceans Canada (DFO) tracks how many fish are caught in commercial fisheries and uses fin-clips or scale samples from captured fishes to provide information on their genetic origins. Coded wire tags, PIT (Passive Integrated Transponder) tags, and acoustic tags provide detailed information on ocean survival, movement, and exploitation rates. These data, and more, help to manage commercial fisheries and are critical to understand, track, and conserve the diversity and distribution of salmon. An evidence-based approach to conserving wild salmon, however, also requires simply counting them. Fortunately for us, and unlike other marine fishes, salmon show up in our proverbial backyards each year for us to count as they return home to spawn (Fig. 1).

**Figure 1.**
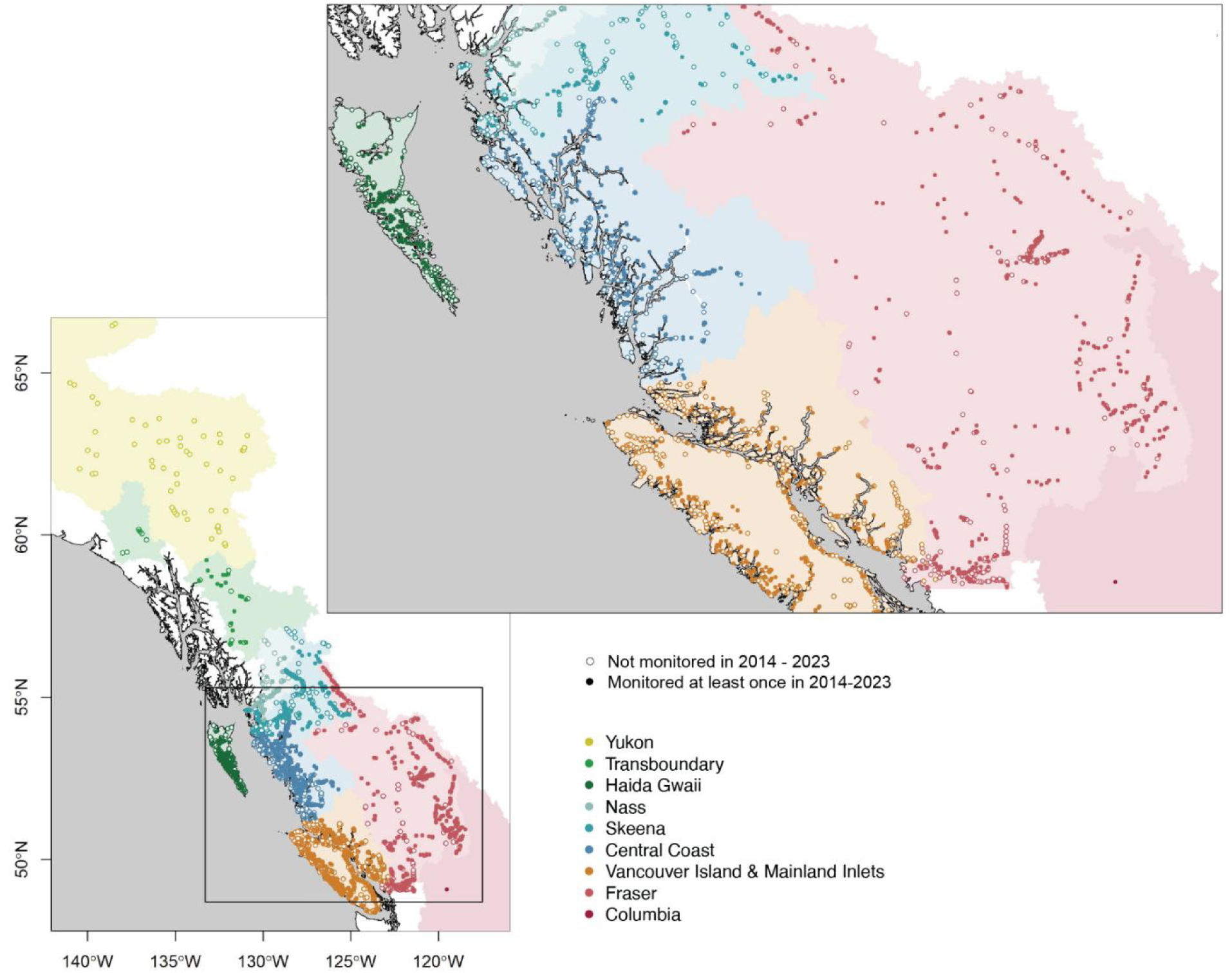
Map of Pacific Canada showing the nine broad-scale regions (colours) and 2,300 sites (points) where spawner monitoring is recorded in NuSEDS. Open circles reflect sites which were not monitored at least once within the 2014-2023 decade.

In Pacific Canada, the history of counting salmon - in one way or another - as they return to spawn extends back thousands of years. For millennia, First Nations communities have developed deep-rooted knowledge of salmon passed down through generations, informed by diverse approaches to monitoring local populations (Atlas et al. 2021a; Reid et al. 2022). This paper focuses on salmon counting programs that extend back to the 1930s, which is relatively modern history but a period for which centrally compiled, broad-scale, and publicly accessible data are available. The evolution of salmon counting coordinated by the federal government over the last one hundred years feeds directly into the data available today. Beginning in the 1920s, the “Annual Reports of Salmon Stream and Spawning Grounds,” commonly known as the “BC16s”, were released annually (Fig. 2). Initially, reports were sent as personal letters to federal fisheries supervisors, describing runs as “good”, “average”, “well seeded”, etc. (McNairnay 1987). In the early 1930s, the reports were formalized to provide an estimated range for the approximate number of spawners on the spawning grounds. This shift was not intended to produce actual estimates of abundance but rather to allow considerations of watershed size, given that “a small stream well seeded could probably have one hundred parent salmon, but a larger stream equally well seeded would have perhaps five hundred parent salmon.” (McNairnay 1987). By the 1980s, the estimates for most populations had transitioned to “point estimates” of the number of individual spawning salmon (Fisheries and Oceans Canada 2023a). In 1995, responsibility for salmon counting moved to DFO’s Science branch, and counting data were brought together in a centralized database in what is now known as the New Salmon Escapement Database (NuSEDS), including details of estimation methods and other descriptive information.

**Figure 2.**
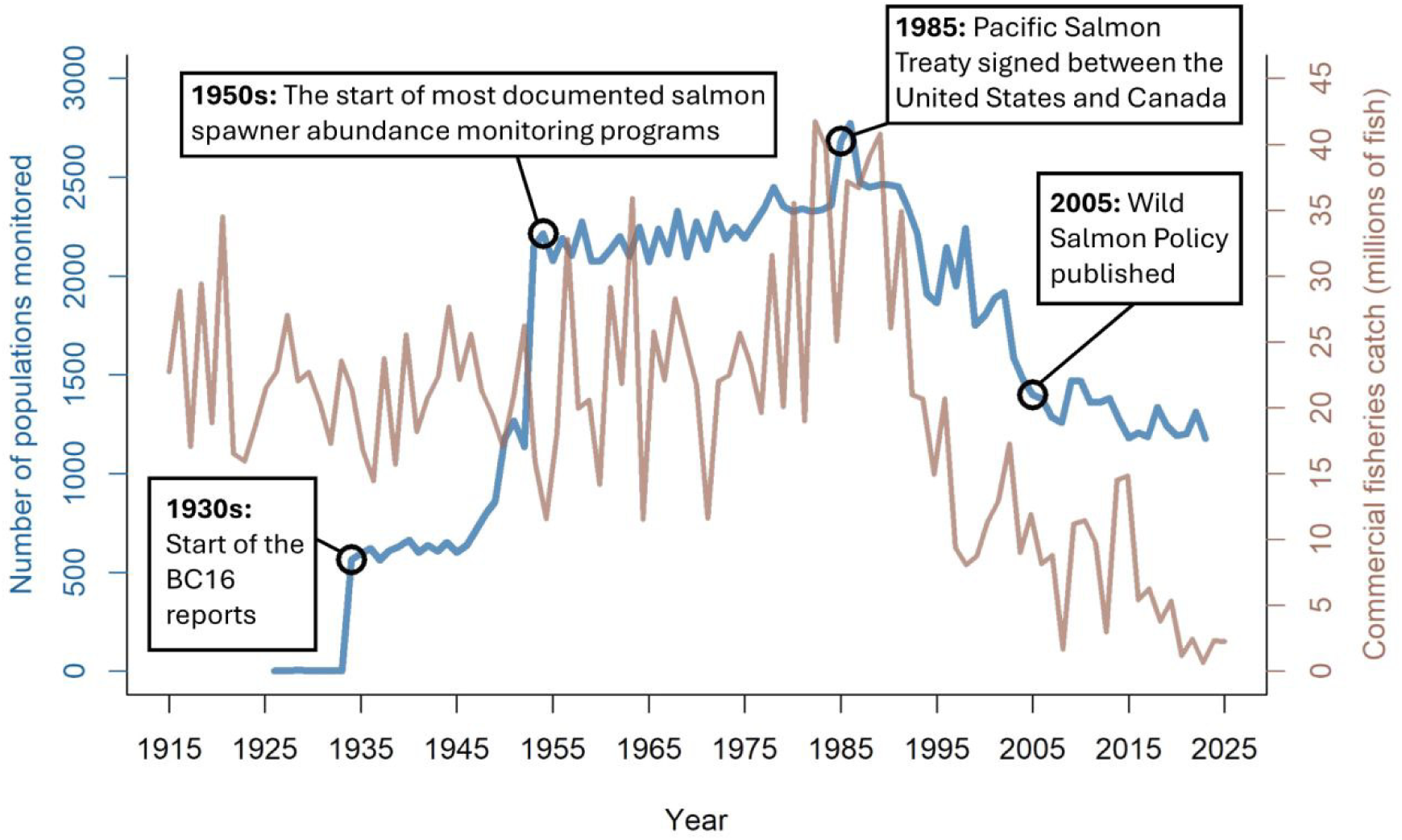
A timeline of the number of populations for which spawner counts appear in NuSEDS (blue) and total commercial salmon catch (brown) in Pacific Canada, highlighting notable events pertaining to abundance monitoring. A population is a grouping of salmon that can return to breed in the same spawning location, usually a stream or river (but sometimes a smaller geographic area). The blue line depicts the number of populations monitored over time (as inferred from the number of records in NuSED, see Methods for full details). The brown line depicts total commercial salmon fishery catch among five salmon species over time as reported by the North Pacific Anadromous Fish Commission (NPAFC 2024). This figure focuses specifically on contemporary salmon abundance monitoring as curated in NuSEDs but many of these populations have been monitored, in one way or another, for thousands of years.

Today, NuSEDS houses the most spatially and temporally comprehensive spawner abundance estimates available for Pacific salmon in Canada. The database is publicly available online and is intended to be updated annually (Fisheries and Oceans Canada 2024a). The specific process for feeding data from local salmon systems to NuSEDS varies by region and data processing protocols are not publicly available. Generally, within a given region, spawner surveys are conducted annually by DFO, local First Nations, consultants, contracted charter patrol officers, and/or local community groups contracted by DFO. The methods for conducting spawner counts vary but generally include stream walks, boat surveys, aerial surveys, snorkel surveys, and fence counts. During the spawning season, regional DFO staff compile and process spawner survey data (e.g., generate total abundance estimates from survey counts using a range of methods). Post-season, DFO NuSEDS staff compile spawner data from across regions and process them for integration into the public database (Fig. 3). In the current Canadian context, NuSEDS is fundamental to assessing salmon status (Box 1), estimating population-level effects of localised stressors (Box 2), and evaluating the efficacy of recovery actions (Box 3).

**Figure 3.**
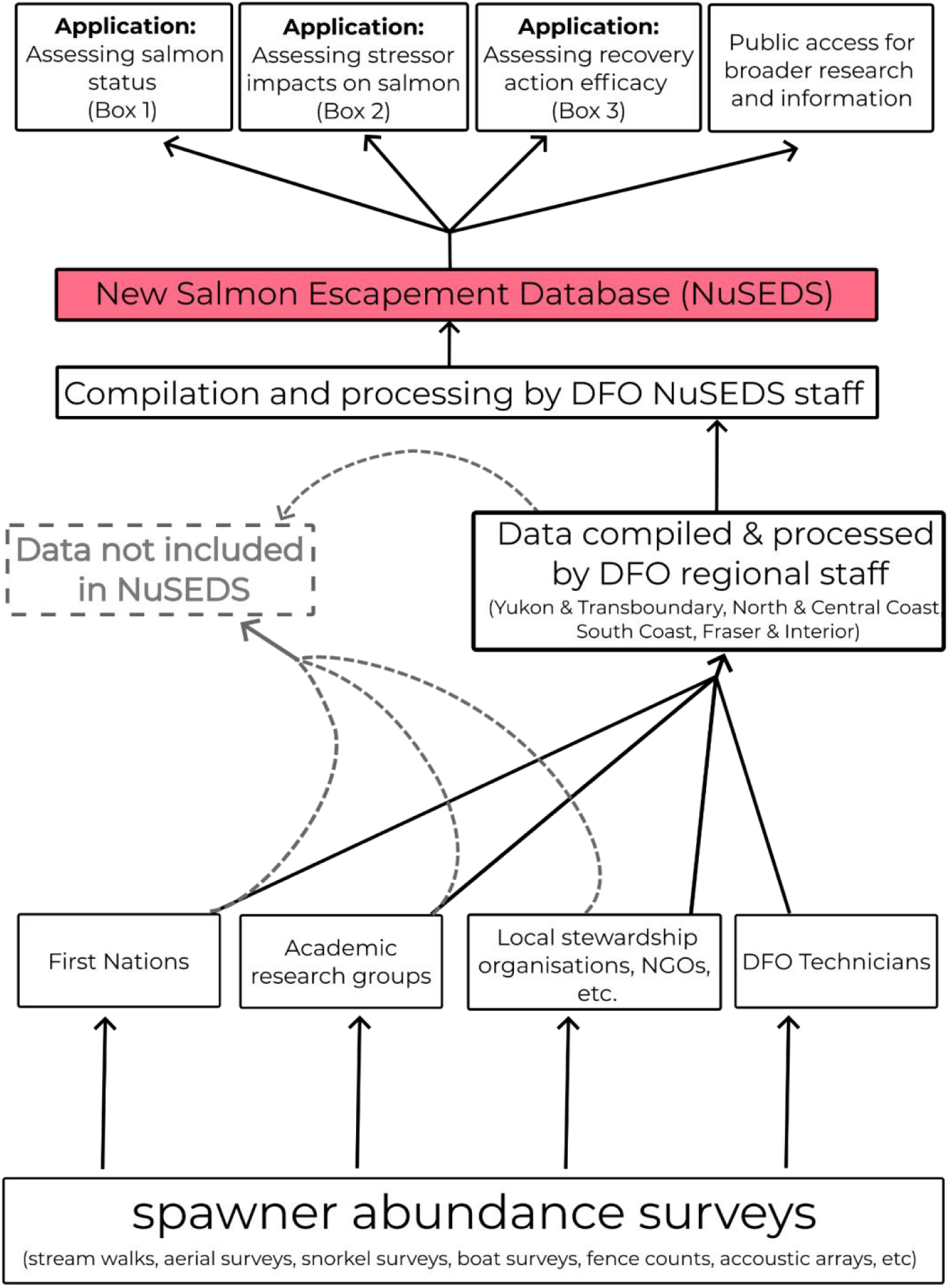
High-level schematic illustrating the data flow and processing steps within the NuSEDS pipeline. General data flow adapted from information provided by the Fisheries & Assessment Data Section at DFO. Spawner abundance surveys are conducted by a range of groups, compiled and processed at a regional level, and subsequently consolidated and processed across regions by NuSEDS staff. Dashed lines highlight places along the data flow where spawner abundance data may be lost from inclusion in NuSEDs.

While rarely acknowledged directly, the importance of counting salmon is implicitly woven into the policy history of Pacific salmon in Canada, most recently embodied in the Wild Salmon Policy and its plan for implementation (Fisheries and Oceans Canada 2005, 2018). The Wild Salmon Policy formalised a shift, from viewing salmon simply as “stocks” to be harvested, toward conservation of the inherent diversity and distribution of wild Pacific salmon. This shift required federal policy to consider what it means to define, monitor, restore, and maintain diverse salmon populations. One step in this process included defining “Conservation Units” (CUs) - genetically and ecologically distinct groups of salmon unlikely to recolonize within a human lifetime if extirpated (Fisheries and Oceans Canada 2005). Delineating CUs at biologically and socially relevant scales is not a trivial task and has been a focus of extensive research (Holtby and Ciruna 2007; Weinstein 2007; Fisheries and Oceans Canada 2009). While imperfect, CUs laid the foundation for implementing the Wild Salmon Policy and, once defined, begged the questions: What data are required to monitor and manage CUs? In what state are those data? Indeed, the first strategy of the Wild Salmon Policy is to “[standardize] monitoring of wild salmon status” (Fisheries and Oceans Canada 2005). For over 20 years, researchers and salmon advocates have noted that the number of populations monitored in various regions of BC has steadily declined (Harvey and MacDuffee 2002; Price et al. 2008, 2017; English 2016; Atkinson et al. 2020; Price and Moore *In review*). However, a coastwide assessment of the trajectory of salmon counting as captured by NuSEDS has not been published. Such an assessment is necessary to understand whether previous calls for increased monitoring have led to change, especially in light of substantial recent investments by the federal and provincial governments through the Pacific Salmon Strategy Initiative ($647 million) and the BC Salmon Restoration and Innovation Fund ($143 million).

Inspired by the late Dr. Jeffrey Hutchings and his commitment to investigating the scientific and policy connections between data and decision-making, we set out to produce an updated, broad-scale assessment of the state of Pacific salmon spawner abundance data in Canada. We focused on NuSEDS, the most comprehensive public database of spawner counts in Pacific Canada and considered monitoring as a two-part process: (1) data collection and (2) data integration into the public database. We assessed the number of populations and CUs monitored annually from 1926-2023 and investigated the relationship between commercial catch, price-per-kilogram, and monitoring effort. Finally, we provide recommendations for optimizing monitoring to meet the conservation objectives laid out in Canada’s Wild Salmon Policy and for improving data management to ensure that timely and accurate information can be shared publicly and appropriately in the coming decades.

## Boxes

### Box #1: Assessing salmon status

The primary metric used to assess biological status of CUs is current spawner abundance (Holt et al. 2009; Pestal et al. 2023; Pacific Salmon Foundation 2024). Estimates of CU-level abundance can be compared to benchmarks - generally set from historical abundances or social-biologically relevant reference points (e.g. the number of fish that maximise fisheries yield) - to determine the status of CUs and determine the potential for fisheries (Chum Technical Committee 2023; Chinook Technical Committee 2024) or the need for conservation intervention. This seemingly simple goal involves hidden complexity and assumptions required to estimate CU-level abundance. In some cases, such as status assessments of geographically constrained lake-type sockeye, spawner abundance can be obtained from a single survey or indicator population that is representative of the aggregate stock or CU. However, in most cases, assumptions must be made to “expand” counts from a limited number of spawner surveys to estimate spawners for an aggregate unit, such as a CU, Stock Management Unit, or region. The expanded estimates are also often required for calculating total returns and exploitation rates, because harvest of salmon does not discriminate between fish that are returning to monitored and unmonitored streams (with the exception of some mark-selective fisheries). Thus, to ensure that catch and spawner data correspond to the same component populations and can be added together, spawner abundance must be extrapolated to unmonitored populations. Although common expansion approaches are fairly robust (Peacock et al. 2020), a severe reduction in the number of monitored populations or decline in monitoring effort over time can invalidate the assumptions of these expansions and create uncertainty around the estimated aggregate spawner abundance. For example, in 2021, only one out of 61 spawning populations was monitored for the West Haida Gwaii chum CU, with a count of 500 spawners (Pacific Salmon Foundation 2023). Based on the historical proportional contribution of the monitored stream to the CU, these 500 spawners were expanded to a CU-level abundance of over 40,000 spawners. Of particular concern is the tendency to stop monitoring salmon populations that have declined to very low abundances and focus on more abundant populations, which may overestimate the expanded number of spawners and present conservation risks to the aggregate on top of potential undocumented extirpations of individual populations (Price et al. 2008).

### Box #2: Assessing impacts on salmon

The spatial and temporal breadth of spawner counts in Pacific Canada has enabled a wide body of research into the effects of local stressors on the responses of Pacific salmon populations. By comparing spawner abundances, or recruits-per-spawner for individual populations, across a range of exposure to a given stressor, the impacts of that stressor on salmon can be estimated. With this approach, research has leveraged salmon abundance data to investigate, for example, the population level-effects of climate change (Hinch and Martins 2011; Malick et al. 2017), parasite transmission from salmon farms (Connors et al. 2010; Krkosek and Hilborn 2011; Peacock et al. 2013), marine mammal predation (Nelson et al. 2019), and forestry (Holtby 1988; Bradford and Irvine 2000; Wilson et al. 2021). All these studies examined stressors that show a range of effect sizes at fine spatial resolution in freshwater or near-shore marine environments (e.g., watersheds or inlets), and many relied on salmon spawner abundance records assembled and maintained by DFO in the New Salmon Escapement Database (NuSEDS) to quantify salmon responses at an appropriate spatial resolution. More spatially constrained data, such as survival estimates from coded wire tags of select indicator stocks, or data with coarser spatial resolution, such as catch within Pacific Fisheries Management Areas, fail to represent the effect sizes and resolution of many pressures. Thus, the decline of spawner abundance monitoring in Pacific Canada limits the ability to quantify effects of localised freshwater and near-shore stressors on salmon abundance and productivity.

### Box #3: Assessing the efficacy of recovery actions

Monitoring spawner abundances before and after recovery actions is essential to track progress towards recovery and facilitate adaptive management. In the past five years, the Canadian government has renewed efforts to address the declining abundance of Pacific salmon by establishing systematic planning processes such as rebuilding plans, prescribed in revisions to the *Fisheries Act,* and Integrated Planning for Salmon Ecosystems, piloted under the Pacific Salmon Strategy Initiative (Fisheries and Oceans Canada 2022, 2024b). For example, resource-intensive efforts led by DFO are underway to create a rebuilding plan for West Coast Vancouver Island Chinook salmon and pilot Integrated Plans for Salmon Ecosystems in the Thompson-Shuswap, Nicola, and Yukon watersheds. While historical spawner abundance data may be sufficient to inform planning and implementation, assessing the success of interventions may be impossible if monitoring declines continue. The efficacy of recovery actions undertaken at finer spatial scales, such as localized habitat restoration or water management interventions, can only be understood by monitoring the affected populations rather than aggregate stocks. Mismatches between recovery strategies and monitoring coverage, both temporally and spatially, may result in instances where significant investments are made in recovery planning and implementation, but the success of such efforts cannot be demonstrated and may be unable to inform future actions.

## Methods

The terminology used for different biological and management scales of salmon is a common source of confusion in analyses such as ours, so we begin with some definitions. For the purpose of our analysis, we define a “population” as the spawners returning to a unique spawning site (e.g., stream, river, or lake) within a CU. These populations mostly reflect the biological unit for which spawner abundances are reported in NuSEDS (Fisheries and Oceans Canada 2024a), except that we consider pink salmon counted at the same site in even- and odd-years as separate populations, consistent with the evidence for genetically distinct even- and odd-year lines (Beacham et al. 2012). A relatively small number (*n* = 87) of NuSEDS populations could not be reliably assigned to a CU; these populations were kept to count the number of populations monitored and were excluded to determine the proportion of CUs monitored (cf. Supplement A). We only included data in our analysis that were available through NuSEDS, and thus define ‘monitoring’ as data collection *and* integration into NuSEDS. In many regions, spawner surveys may be undertaken but the data are either not submitted to or not included in NuSEDS. This potential reporting bias is addressed in recommendation #4 in the Discussion.

We downloaded two datasets from NuSEDS on January 21st, 2025 (Fisheries and Oceans Canada 2024a). The first dataset (*all_areas_nuseds*) contains the annual counts (e.g. number of natural adult and jack spawners) for each population. The second dataset (*conservation_unit_system_sites*) associates each population to its respective CU and site. We merged the two datasets to associate each population’s time series to its CU. There were numerous errors and omissions between these two datasets including duplicate time series, abundance estimates assigned to wrong populations or CUs, or missing CU information. We undertook a systematic review of the data to identify and correct these cases (cf. Supplement A for details and access to R code). The resulting cleaned spawner survey data are available online (Carturan and Peacock 2024).

We assigned each population to one of nine broad-scale regions based on the site location (latitude/longitude) in NuSEDS to assess broad spatial patterns in monitoring (Fig. 1). The final dataset contained spawner abundance data from 1926 to 2023 for the five Pacific salmon species (Chinook, chum, coho, pink, and sockeye), 392 CUs, 6,973 populations, and 2,345 sites (Table 1).

**Table 1.** Number of Conservation Units (CUs), populations, and sites present in the NuSEDS data in each region after cleaning (zeros and NAs counts were removed). Sites are unique locations where spawners from one or more CUs may be counted. Populations are unique combinations of CUs and sites.

| Region | Number of CUs | Number of populations | Number of sites |
| --- | --- | --- | --- |
| Yukon | 15 | 60 | 54 |
| Transboundary | 22 | 65 | 37 |
| Haida Gwaii | 28 | 756 | 233 |
| Nass | 20 | 307 | 82 |
| Skeena | 54 | 777 | 265 |
| Central Coast | 111 | 1681 | 404 |
| Vancouver Island & Mainland Inlets | 83 | 2297 | 666 |
| Fraser | 57 | 1028 | 603 |
| Columbia | 2 | 2 | 1 |
| **Total** | **392** | **6,973** | **2,345** |

From the cleaned NuSEDS data, we calculated the annual “number of populations monitored” as the number of populations having a numeric estimate in any of the following fields: natural adult spawners, natural jack spawners, total natural spawners, adult broodstock removals, jack broodstock removals, total broodstock removals, other removals, or total return to river. There were 3,449 yearly counts (2%) with a maximum value of zero across these fields. We found abrupt increases in the frequency of zeroes starting in certain years and only for certain regions and species (e.g., Fig. S6). Given the available information, it was not possible to determine whether these zeros represented monitored populations with true zero counts. As such, we opted to conduct the analysis under two different assumptions: in the main text we present results assuming that counts of zero indicate no monitoring and in Supplement B, we present results under the assumption that monitoring did occur when zero spawners are reported.

We divided the number of populations monitored each year by the total number of populations monitored to obtain the annual “proportion of populations monitored”. The annual “proportion of CUs monitored” was the number of CUs with at least one population monitored divided by the total number of CUs (which differed in even and odd years because pink salmon CUs are only monitored every other year). We summarised declines in monitoring for each species and region as the average change per year in the number of populations monitored. We estimated the average change per year as the slope of a linear regression model fitted to the number of populations monitored each year from the peak of monitoring in 1986 to 2023. Spawner monitoring arises, in part, from the need to make informed commercial fishery management decisions, such as defining Total Allowable Catch, estimating exploitation rates, etc. If commercial fisheries are a significant driver for monitoring, reducing the size, value, and/or spatial extent of these fisheries might be expected to result in corresponding declines in monitoring given lower urgency or fewer requirements to generate quantitative estimates that rely on abundance data. To investigate whether and for which species this relationship existed, we quantified the association between the number of populations surveyed and commercial catch (kilograms), price per kilogram, and species. We defined two models to compare the effect of landed value (the product of catch and price) and catch alone. We fitted a negative binomial generalised linear model (GLM) with the log-link function and compared the two models using AIC and pseudo-R^2^. Further details of model fitting and assessment can be found in Supplement B. We downloaded catch data from the North Pacific Anadromous Fish Commission website (NPAFC; 2024) in July 2024. We considered the Canadian, commercial catches expressed in kilograms for each of the five Pacific salmon species. We imported the price-per-kg for each species from DFO’s Economic Analysis (Fisheries and Oceans Canada 2023).

We used R (version 4.5.0; R Core Team 2025) and the R packages `tidyr` 1.3.1 (Wickham et al. 2024) and `dplyr` 1.1.4 (Wickham et al. 2023) to handle and analyse the datasets and produce the figures. We used the R packages `sf` 1.0-20 (Pebesma and Bivand 2023) to group sites in their respective region, `stats` 4.5.0 to fit the GLMs. The workflow is fully reproducible by accessing the R scripts and datasets in the Zenodo repository at https://zenodo.org/records/14248905.

## Findings

Monitoring and reporting of salmon spawner abundances throughout BC and the Yukon peaked in the mid-1980s, with over 2,700 individual populations across ∼1,300 unique sites being monitored annually (Fig. 2). Since 1986, there has been a marked decline in the number of populations monitored annually, with an average loss of 41 populations per year (1.5% per year) in the NuSEDS data from 1986 to 2023. Across 6,973 unique populations (Table 1), only 37% have spawner records in the most recent decade (2014-2023) compared to 69% in the period 1980-1989 (i.e., a decline of 31%).

Coho and chum populations saw the greatest decline in monitoring, particularly for Vancouver Island & Mainland Inlets, the Central Coast, and Haida Gwaii (Fig. 4c,f,g, Fig. S1b,c). Pink salmon also saw major reductions in monitoring on the Central Coast, Skeena, and Haida Gwaii (Fig. 4c,f,e, Fig. S1d). Fraser Chinook and Fraser sockeye saw the only increases in monitoring since the 1980s among all regions and species (Fig. 5), although the number of Fraser Chinook populations monitored peaked later, in 2000, and has also since declined (Fig. 4h).

**Figure 4.**
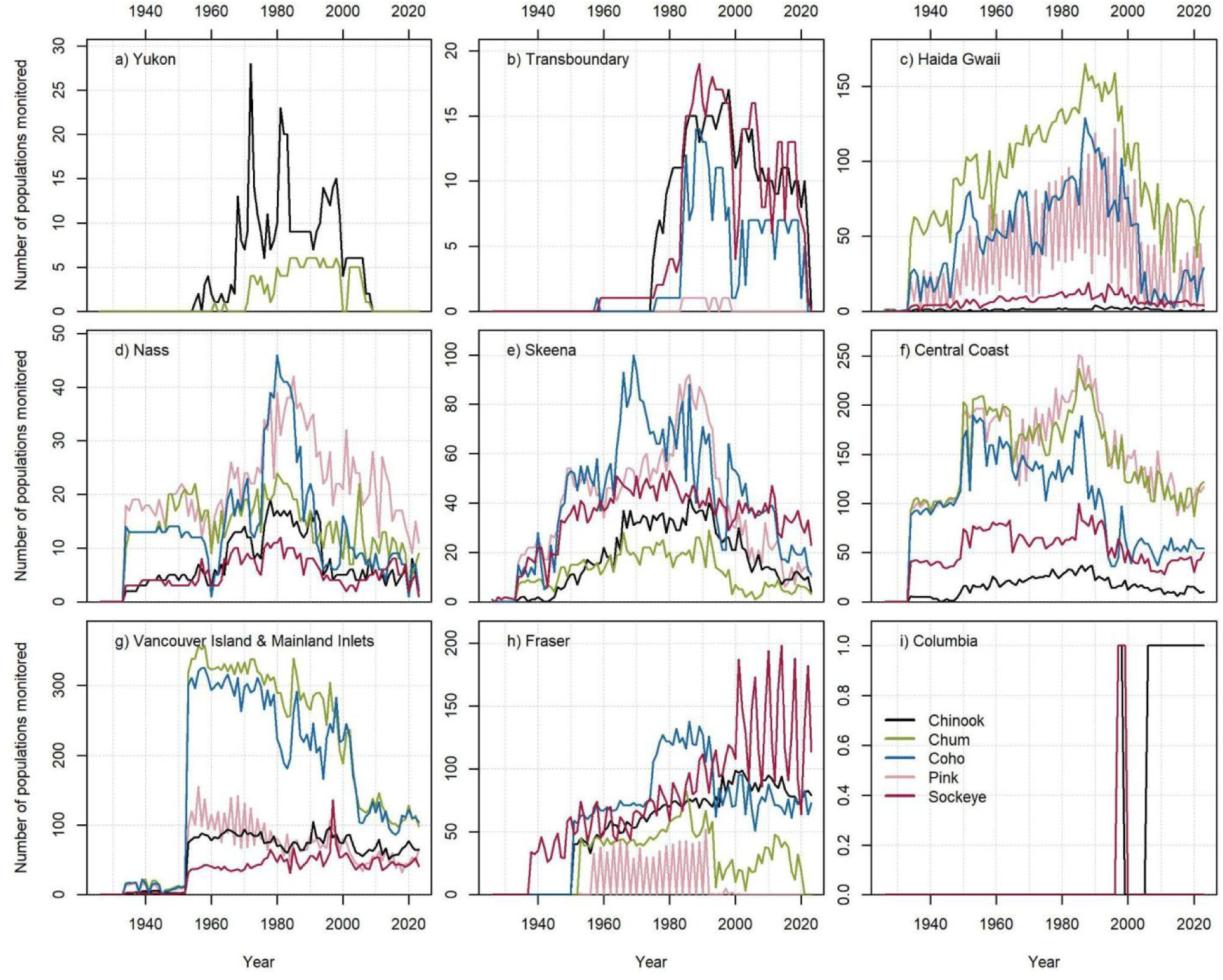
The annual number of salmon populations monitored in NuSEDS, broken down by region (panels a through i) and by species (Chinook, chum, coho, pink, and sockeye).

**Figure 5.**
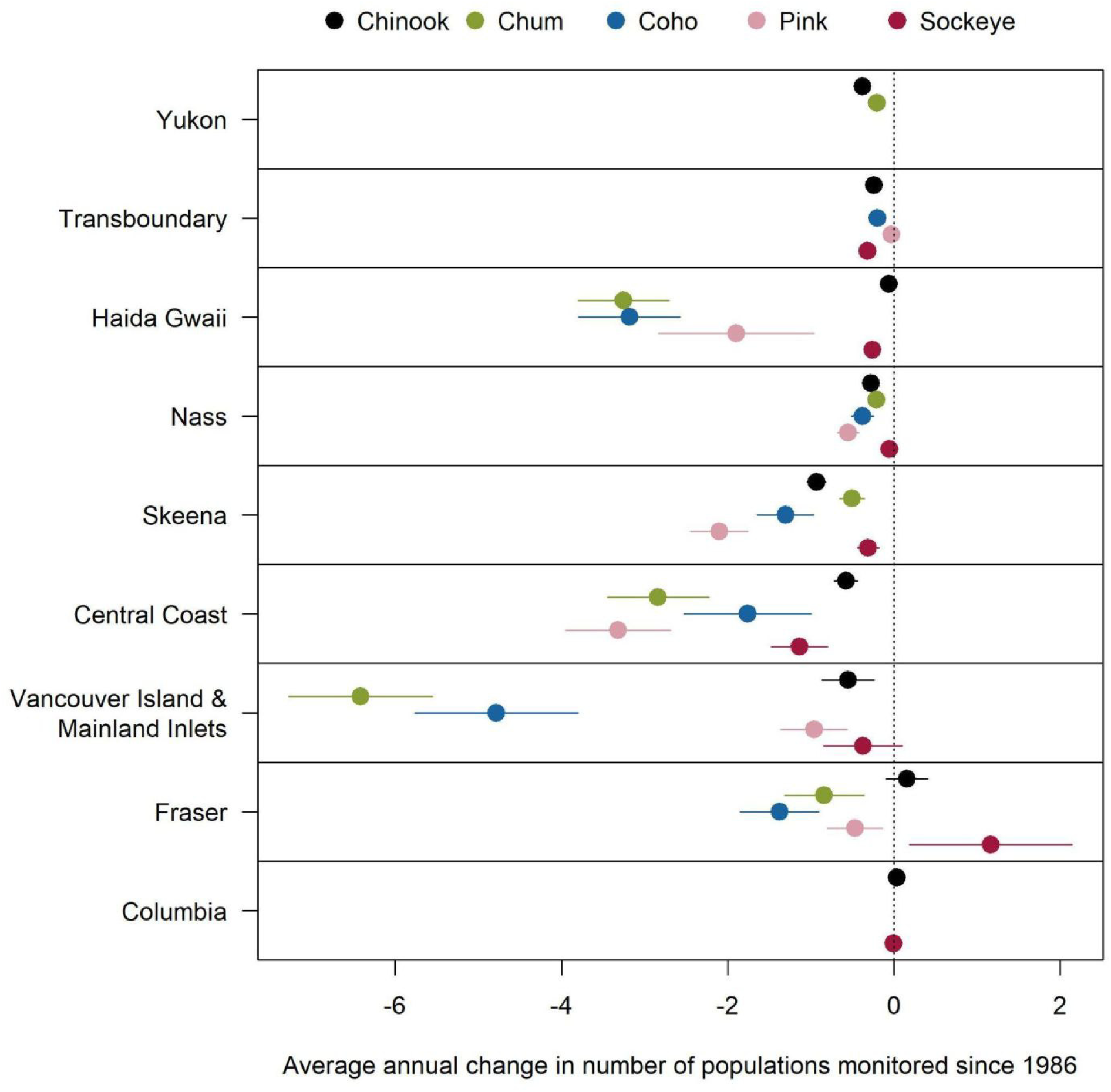
The average annual change in populations monitored since 1986 as estimated via linear regression for each species and region. Dots and horizontal bars represent the slopes and their 95% confidence intervals, respectively.

Trends in monitoring effort across species and regions through time tend to reflect trends in commercial catches, particularly over the last 40 years (Fig. 2). For example, declines in coho monitoring in the early 2000s coincided with the moratorium on coho fisheries following a period of low marine survival that started in the 1990s (Decker et al. 2014). When there was no longer such a strong need to inform coho fishery management, monitoring diminished. Additionally, this reduction in coho monitoring may have inadvertently led to a parallel decline in chum monitoring, most notably in the Vancouver Island & Mainland Inlets region (Fig 3g). Given their overlapping spawn timing, chum may have been monitored opportunistically during surveys targeting coho. Despite the increase of many coho populations in the late 2000s and 2010s, with the reduction of harvest and more recent shift in ocean condition (Wilson et al. 2025), monitoring has not returned to historical coverage levels. As the Interior Fraser coho Management Unit recently exceeded historical average abundances for the first time in decades (Connors et al. 2024), we anticipate the resumption of fisheries. Whether the diversity and distribution of coho within the Fraser watershed have also recovered, and how vulnerable this stock is to another collapse, are difficult to assess in the absence of broad-scale river-level monitoring (Boxes #2-3). Pervasive declines of chum salmon over the past five years (North Pacific Anadromous Fish Commission 2023; Connors et al. 2024) have added to the urgent need to better understand the abundance and distribution of these two species in late-season spawner surveys.

We found some empirical support for the hypothesis that monitoring effort reflects economically important fisheries. There is a positive relationship between the number of monitored populations and the annual landed commercial value for coho, chum and pink (Fig. 6). The model with landed value as a predictor was better supported than the model with catch (Δ AIC=30.2). Overall, there is a strong temporal component to this relationship, with more recent years having low monitoring effort and landed value. Chinook and sockeye show the lowest levels of association among species between monitoring effort and landed value, a pattern reflecting the relatively low monitoring of these species in the early part of the time series when catches were moderate, and relatively high levels of monitoring as catch declined in the 1990s and 2000s. Several general and contextual factors may explain the discrepancy observed for the two species. First, both species hold high social and cultural value independent of their economic value and may also be more frequently subject to monitoring requirements through treaty obligations, species-at-risk processes, etc. Second, the relationship for sockeye might be influenced by the significant decline in Fraser Sockeye returns during the 1990s that prompted investigations into potential causes and the formulation of recommendations (e.g., Pacific Fisheries Resource Conservation Council 2002; Lapointe et al. 2003; Cohen 2012). This notable event may explain why Fraser Sockeye is the only species-region combination where monitoring effort has increased since the late 1990s (Fig. 4). In the other regions, however, the total number of monitored sockeye populations have experienced declines and the overall proportion of monitored sockeye CUs in Pacific Canada has declined (Fig. 7c). Third, recreational fisheries have been the dominant source of Chinook harvest at least since 2000 - a trend unique to this species (Fisheries and Oceans Canada 2023). Consequently, commercial landed value might not represent the species’ economic value accurately.

**Figure 6.**
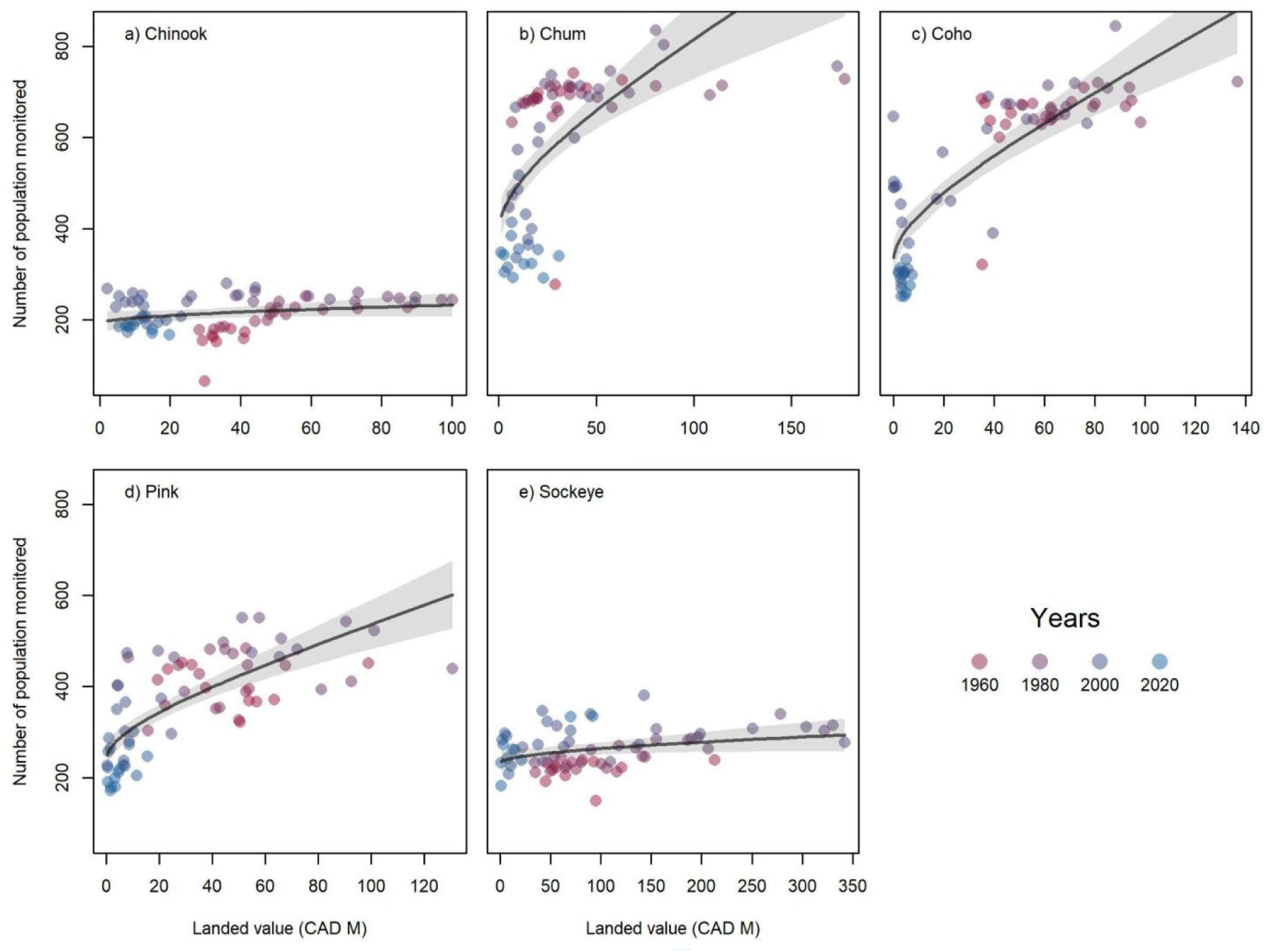
Prediction of the number of populations monitored as a function of landed value (in million of CAD), which is the product between commercial catch (in kg) and price (in CAD per kg). A generalized linear model with a negative binomial distribution and log-link function was fitted with catch, price and species as predictive variables (n = 345). Lines and polygons represent model fits and 95% confidence intervals, respectively; circles represent the data, which span from 1952 and 2020. Zero counts in NuSEDS were filtered out.

**Figure 7.**
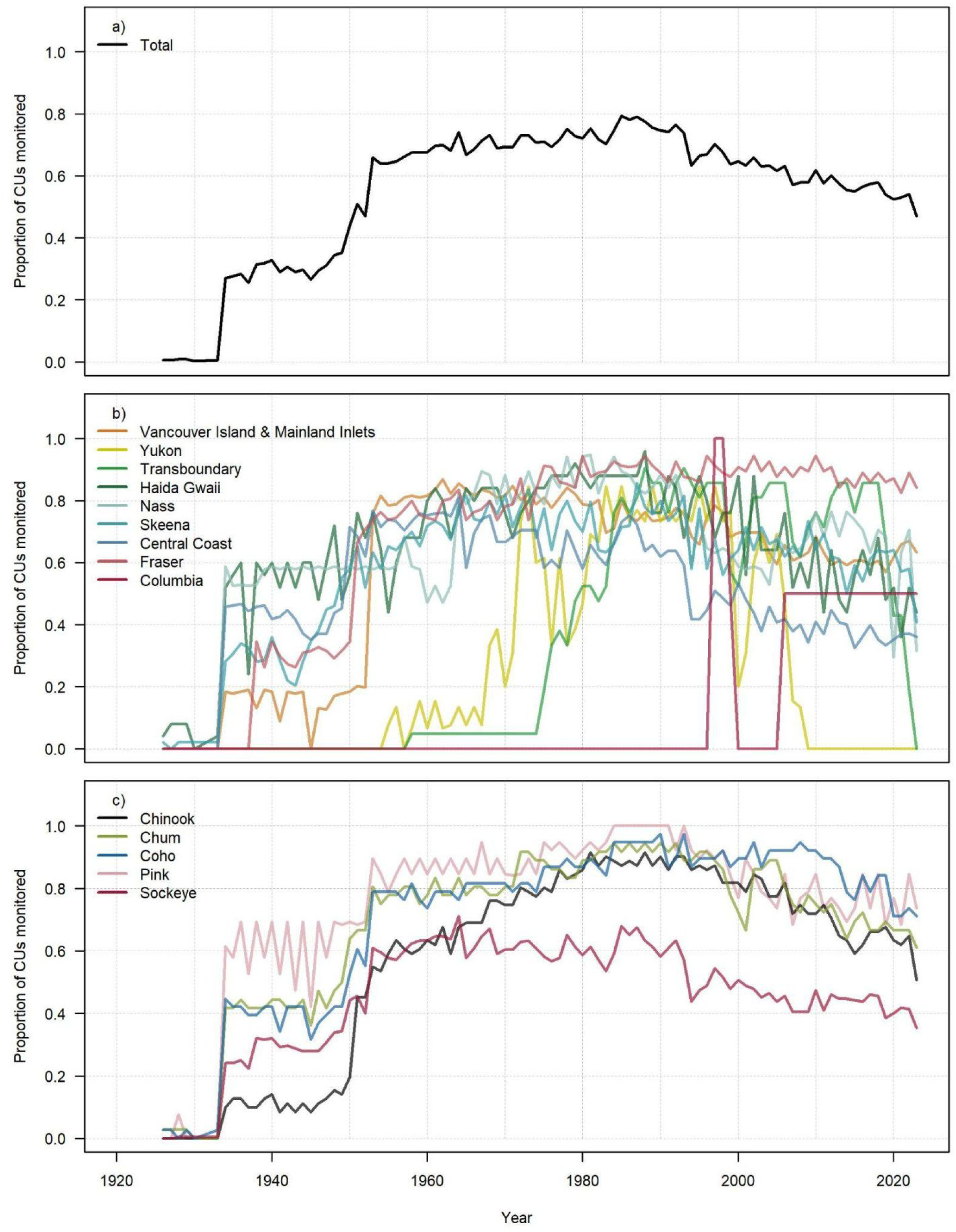
Annual proportion of conservation units (CUs) with at least one population monitored in NuSEDS.

Evidence suggests that the value of commercial fisheries influences spawner monitoring effort, but multiple factors likely contribute to the history of spawner monitoring over time. We cannot account for every nuance that drives decisions regarding monitoring effort but, with limited resources, high-profile populations tend to be prioritised. But this approach to prioritisation does not implicitly guarantee optimal monitoring with respect to meeting conservation objectives such as those outlined in the Wild Salmon Policy. In an example of this pattern, 14 of the 30 most frequently monitored Chinook populations in the Vancouver Island & Mainland Inlets region are “indicator stocks” reported on by the PSC’s Chinook Technical Committee, with an average of 64 years of data per indicator population. By contrast, the 255 Chinook populations in the region that are not indicator stocks have an average of 17 years of data, and 66% of them have not been monitored at all in the most recent decade (2014-2023).

Even within streams, monitoring tends to increase in years with more abundant brood lines. As a result, pink salmon are often predominantly monitored in either even or odd years (e.g., Fig. 4b-h). In the Vancouver Island & Mainland Inlets region, monitoring was historically predominant in even years, following the dominant return year (Fig. S3e,f). “Cyclic” sockeye in the Fraser River, which show marked patterns across four brood lines, tend to have higher monitoring effort in the higher-abundance “dominant” years, a pattern that has intensified since the early 2000s (Fig. 4h). While this prioritisation may have practical justifications, it also makes it difficult to detect shifts in dominance patterns - which have already been shown to occur in some pink salmon populations (Krkosek et al. 2011; Irvine et al. 2014) - and declines in non-dominant years.

A corollary of “monitoring for fisheries” is the dearth of data for less abundant populations. These monitoring data are, however, essential for understanding patterns of salmon distribution and diversity, which include both large and small runs. Monitoring only abundant populations reduces the power to detect shifts in abundance, such as shifts in dominance for pink salmon (Irvine et al. 2014), or declines in biodiversity, such as the erosion of sub-dominant and off-cycle lines for Fraser sockeye (Ricker 1997), that may affect the sustainability of management decisions and the resilience of salmon. Furthermore, ground-based salmon counting surveys provide valuable information beyond abundance counts. They are an opportunity to monitor the system more broadly, including documenting disease events, pre-spawn mortality, river blockages, and high temperatures. Less abundant or declining populations have been termed “ghost streams” in reference to their undetermined status and potential for extirpations to go unrecorded (Price et al. 2008).

The proportion of CUs with at least one population monitored has also broadly declined since the mid-1980s across all regions and species (Fig. 7). CUs were only defined after the implementation of Canada’s Wild Salmon Policy in 2005 (Fisheries and Oceans Canada 2005), which emphasized the importance of monitoring these ecologically and genetically unique groups of populations. Despite this recognition, monitoring of CUs has declined since 2005 (Price et al. 2017; Price and Moore *In review*), continuing even after 2018 when DFO renewed its commitment to the policy with the release of the Wild Salmon Policy Implementation Plan (Fisheries and Oceans Canada 2018).

In some regions, patterns in monitoring may reflect a reporting bias rather than an actual reduction in effort, although it is difficult to disentangle the two if the data are not made available. For example, no data have been included in NuSEDS for Yukon populations since 2008 (Fig. 4a), yet spawner surveys are reported in other publications (Yukon River Joint Technical Committee 2025a). In other cases, the apparent disappearance of monitoring may reflect a shift towards aggregate assessments at more downstream locations. This is the case for Fraser River pink salmon, for which monitoring of individual spawning populations appears to have stopped in the 1990s (Fig. 4h) when the Pacific Salmon Commission began recording pink salmon abundance using a hydroacoustic station near the mouth of the Fraser (Glaser et al. 2025). Although this method provides rapid and accurate counts of the number of pink salmon returning in-season to the Fraser, and is used to inform fisheries management, the population-level survival of those fish to their spawning grounds and distribution throughout the Fraser River basin is now unknown.

We define monitoring effort as the number of populations monitored, without accounting for differences in data quality, which may be a limitation. Indeed, higher quality spawner surveys (e.g., sonar and video weirs) are more expensive and could therefore be considered to represent higher monitoring effort than simpler surveys (e.g., stream walks). Since these surveys were more likely implemented in the later part of the assessed time period, the decline in the number of populations monitored might reflect, to some extent, the trade-off between quantity and quality (i.e., higher quality abundance estimates from fewer systems). Although data quality is reported in NuSEDS, we excluded it from our analysis because consistent data quality scores are only available from 1998 onward (Fig. S7). However, declines in the number of populations monitored are evident across all survey quality categories over the available time period (Fig. S9). This suggests that the reduction in monitored populations is unlikely to be the result of a strategic shift toward higher-quality surveys focused on fewer populations, but rather reflects a broader decline in overall monitoring investment.

Retaining observations with zero counts in the NuSEDS data yields similar results overall (Table S1; Figs. S4, S8, S10, S22). The main differences pertain to sockeye populations in the Fraser, which accounts for 56% of the zero counts, most recorded after 1998 (Fig. S6). This abrupt reporting of zero counts dampens the magnitude of fluctuations in monitoring effort after 2000 (Fig. S4h) and increases the average annual change in monitored population since 1998 (Fig. S5). While these differences do not undermine our conclusions, they underscore the extent to which the interpretation of results hinges on the consistency and reliability of how zero counts versus NAs are reported.

## Discussion

Spawner abundance monitoring and reporting has historically focused on informing commercial fisheries management, resulting in monitoring efforts being concentrated on economically important species and numerically abundant populations. Commercial catches of salmon in BC and the Yukon have collapsed in recent years (Walters et al. 2019; Yukon River Joint Technical Committee 2025), along with the effort towards collecting and sharing spawner data that was previously used to inform management of those fisheries. For the commercial salmon fisheries that persist, lack of spawner abundance data has led to loss of sustainability certifications (Raincoast Conservation Foundation 2019; Marine Stewardship Council 2022) which impedes access to diverse markets that increasingly favour sustainable seafood. While well-managed salmon fisheries will continue to be sustainable into the future, the current magnitude of commercial fisheries is very different than it once was, and monitoring priorities must shift to reflect the changing seascape of salmon fisheries. As public interest and funding shifts towards salmon conservation and recovery planning, the limitations of past, commercial fishery-oriented monitoring objectives are becoming apparent: we currently lack the data to assess the status of diverse Conservation Units or understand threats to their survival (Boxes #1-2). The absence of this information impedes recovery planning and the assessment of recovery actions (Box #3).

We recognize that the patterns we observed may be due to a combination of declining monitoring effort and potential reporting bias. It takes time and effort for data to be verified and entered into NuSEDS, creating the possibility of lag that may be misinterpreted as a decline in effort. However, the long time period over which we observed consistent declines in monitoring - more than 40 years - suggests that such a lag is not the only factor at play. We observed some regions and species where monitoring appears to have declined or completely stopped based on the NuSEDS data even though we know that spawner surveys have continued and are reported elsewhere. For example, in the Yukon and Transboundary regions, many spawner surveys are reported on by the Technical Committees tasked with informing salmon management under the Pacific Salmon Treaty (Transboundary Technical Committee 2022; Yukon River Joint Technical Committee 2025b), but the very same surveys are inconsistently updated in NuSEDS (Peacock et al. *In press*). Regardless of whether data continue to be collected, their usefulness to research and broad-scale prioritization and planning is undermined if they are not compiled and included in a single, centralized, and publicly accessible database - NuSEDS.

As climate change requires salmon to adapt, tracking changes in their distribution and diversity through spawner surveys has never been more critical. Monitoring programs that rely on a handful of key indicator stocks may miss potential changes in diversity, which underpins the resilience of salmon social-ecological systems (Schindler et al. 2010; Nesbitt and Moore 2016). Numerically abundant indicator populations may persist as surrounding (unmonitored) populations decline, and aggregate stocks may continue to have large returns even as the diversity of their components is undermined (Price et al. 2021). This can lead to apparently healthy populations, until a tipping point is reached or a catastrophic event threatens the viability of the aggregate stock. For example, preceding the collapse of Atlantic cod fisheries in eastern Canada, aggregate offshore surveys continued to suggest healthy cod stocks, while inshore fishers observed local extirpations - proverbial canaries in the coal mine (Hutchinson 2008). Climate changes are now exacerbating other pressures on salmon, yet insufficient data hinders our ability to understand the adaptive capacity - or fragility - of salmon populations. Here, we provide recommendations with the goal to encourage and enable strategic investments in spawner abundance monitoring that can support informed, evidence-based decisions about salmon conservation, management, and recovery.

## Recommendations

We, the authors, are non-Indigenous researchers with extensive technical experience working with large salmon datasets to serve the scientific objectives of a broad range of groups including academic researchers, non-governmental organisations, federal and provincial governments, First Nations governments, and other Indigenous organizations. We are best positioned to make recommendations that would improve the technical utility of salmon abundance data generally and NuSEDS in particular. We also recognize that data (and their handling) are intertwined with the decisions they inform, and we draw on our experience and a growing body of literature on Indigenous Data Sovereignty to offer suggestions intended to be respectful of Indigenous rights.

Solutions to the evident decline in monitoring will involve increased investments in boots on the ground, but the burden of the costs can be mediated by strategic investments in collecting data, employing new technologies, and making improvements to how data are processed and shared.

### Recommendation #1: Optimize spawner monitoring to serve the assessment of salmon diversity, for the purpose of maintaining it

In the Wild Salmon Policy (Fisheries and Oceans Canada 2005, 2018), DFO identified monitoring wild salmon status as the foundation of its strategy to accomplish a core objective: “restore and maintain healthy and diverse salmon populations”. However, our assessment of current monitoring efforts suggests that the institutional shift required to apply this strategy has yet to occur (also, see Price and Moore *In review*). We have shown that, historically, investments in spawner surveys have mirrored commercial catches through time. As commercial catches have declined to zero in some regions, accompanying declines in monitoring efforts have undermined our ability to assess salmon status, understand threats, and implement effective recovery plans. Clarifying monitoring objectives in-practice to include assessing changes in salmon biodiversity would help inform conservation and recovery efforts focused on maintaining that diversity, while promoting sustainable management of modern-day fisheries - balancing local and regional objectives, First Nations rights, and both recreational and commercial interests.

Annual spawner surveys of all ∼2,345 locations listed in NuSEDS (Table 1) may not be feasible, but strategic monitoring programs can be designed to meet clearly stated objectives with minimal costs, making the most of limited resources. Spawner monitoring focused on assessing the status of Conservation Units, as outlined in the Wild Salmon Policy, provides a starting point for assessing and conserving salmon diversity that is necessary to support sustainable fisheries and ensure salmon resilience in the face of environmental change. With almost 400 individual CUs (Baxter and Hamilton 2024), many of which are remote and logistically challenging to monitor, annual monitoring of each may not be possible. When prioritisation among CUs is necessary, monitoring could aim to distribute effort across all salmon-bearing regions rather than focus on the regions with the most high-profile populations or historically-renowned fisheries.

Some CUs encompass hundreds of spawning populations and thus monitoring a single site is likely insufficient. In such cases, the total number of populations of interest may exceed the capacity for monitoring. Answering the simple question, ‘how many populations should we count?’ is deceptively difficult because it depends on the monitoring objectives, the nuances of particular regions, and the available resources. While it is beyond the scope of this paper to provide specific numbers, an interested reader can refer to abundant resources on the design and implementation of salmon monitoring programs (National Centre for Ecological Analysis and Synthesis 2010). Extensive work has been published on regionally specific monitoring designs (English 2016; Atlas et al. 2021b), quantifying consequences of missing spawner data in run reconstructions, and optimal monitoring designs to assess conservation status (Holt et al. 2011; Peacock and Holt 2012). Techniques such as “rotating panel” monitoring designs (Peacock and Holt 2012) can be employed to maximize the information obtained with limited resources. These designs, where some indicator populations are assessed annually while other populations are assessed on a rotating basis, allow one to efficiently detect contractions in the spatial distributions of spawners.

### Recommendation #2: Provide sufficient and consistent resources to spawner monitoring programs to meet stated objectives

The Wild Salmon Policy set out a forward-looking roadmap for the conservation of wild Pacific salmon that included monitoring and assessment of Conservation Units, but the challenge of following that roadmap has not been met with sufficient and consistently available resources. Ironically, there have recently been historic investments in salmon ($647 million) through the Pacific Salmon Strategy Initiative (Government of Canada 2021) and BC Salmon Restoration and Innovation Fund (Government of Canada et al. 2019). We argue that these investments, while unprecedented in their total amount, have not led to the stable long-term funding and coordinated approach needed to rebuild salmon monitoring in Pacific Canada.

A major barrier to revitalizing spawner monitoring in Pacific Canada is the apparent lack of integration and communication between different initiatives within DFO. Rebuilding spawner monitoring requires long-term funding but it will also require a coordinated approach to identify short- and long-term priorities, streamline data handling processes, and coordinate within and between regions. There are likely multiple ways to meet this challenge, but we suggest two specific targets for reinvestment at each level of the spawner monitoring data flow (Fig. 2). First, we recommend investing to build capacity within the NuSEDS team to support their work developing and maintaining a quality-controlled, well-documented database (see Recommendation #4). Second, we recommend investing in well-staffed regional DFO teams with capacity to steward spawner data from collection to integration within NuSEDS and support the many non-DFO groups conducting spawner abundance surveys. Systems with accountability for collecting, documenting, and making information accessible must be a core responsibility for all DFO staff tasked with managing data.

Investment to reinvigorate spawner monitoring across BC and the Yukon is a common-sense approach to salmon conservation and management. It is also a practical opportunity to create local jobs in the many rural coastal and inland communities working to expand and diversify labour opportunities. Stream-walking jobs to monitor spawning salmon are community based, provide connection between people and places, and have the potential to be purposeful, meaningful occupations for people in Pacific Canada. Reviving salmon counting will likely involve reinvesting in DFO-led stock assessment programs, but in many salmon watersheds the practical solution may involve supporting local communities and First Nations already stewarding watersheds and salmon. For example, DFO reported working with 40 unique Indigenous collaborators in 2023 and 2024 on salmon stock assessment activities (Fisheries and Oceans Canada 2024c). Many locally based salmon counters are funded through DFO contracts that have shrunk in value and scope year-over-year (e.g., Billy Proctor, *personal communication*, 2022). Nonetheless, these groups often have ample expertise and enthusiasm, and sufficient investment can enable them to count more populations.

### Recommendation #3: Invest in new technologies that can improve monitoring efficiency

The rapid advancement of computer analysis and remote sensing has significant potential to improve the quality of data and reduce costs. Historically, new monitoring technologies have focused on providing more in-depth knowledge about survival of particular stocks, rather than improving broad-scale coverage of monitoring. For example, coded wire tags, PIT tags, and acoustic tags provide detailed information on ocean survival and movements but are only deployed on a limited number of stocks, requiring strong assumptions (e.g. about the representativeness of coded wire tag stocks) to extrapolate findings to broader scales. Emerging technologies, on the other hand, may make monitoring more efficient, thereby allowing for increased coverage. The application of computer vision and deep learning to process salmon sonar and video data, collected at in-river weirs, can reduce the need for manual review of hundreds of hours of footage, with potential to reduce costs significantly (Atlas et al. 2023). Uncrewed aerial vehicles (UAVs), commonly known as drones, have been shown - in amenable systems - to provide reliable indices of salmon spawner abundance for relatively low cost (Ponsioen et al. 2023). Environmental DNA, or eDNA, is another emerging technology that may be suitable for detecting, and potentially estimating abundance of, salmonids (Levi et al. 2019).

Although these technologies are in their infancy, and cost savings may not be immediate, investing in their development and application in Canada could speed up broad-scale application and its associated benefits. To ensure salmon abundance data maintain their value as long time series, it will be critical that new technologies are applied together with existing monitoring approaches to validate and calibrate between methods.

### Recommendation #4: Compile and maintain high-quality spawner data across government regional divisions and organizations

In some cases, monitoring gaps may be due to a failure to integrate existing spawner abundance data into NuSEDS, a process which is slow or inconsistent across regions. NuSEDS represents the most comprehensive dataset on salmon abundance, and yet we found many instances where data were not contributed to NuSEDS despite being published by DFO in other reports or made available on request. We mentioned examples in the Yukon and Transboundary regions above, but the monitoring gaps for Columbia sockeye and Chinook salmon (Fig. 4i) can also be filled by looking at alternative sources (Hyatt and Stockwell 2019; Matylewich et al. 2019). For Fraser chum salmon, the most recent data in NuSEDS are from 2020, four years behind collection, and the last year of data currently available for the Harrison River (a major chum population of the Fraser watershed) is 2015, almost 10 years ago. This lack of data centralisation limits the power and application of the data to improve our understanding and timely management of Pacific salmon (e.g. Boxes #1-3). In the context of this study, it also hinders our ability to document putative declines in monitoring effort. Following the collapse of Atlantic cod and declines in Pacific salmon, several noted experts recommended that all scientific information on stock abundances be made public as soon as it is presented to DFO (Hutchings et al. 1997). Even ‘interim’ or ‘provisional’ data are valuable for timely detection of conservation risks and should be made available. Any errors could be corrected in future versions of the data. As an excellent example of this proactive approach, the Pacific Salmon Commission Secretariat releases ‘in-season’ estimates of Fraser sockeye abundance that are retroactively finalized and re-released pending estimates of pre-spawn mortality (Pacific Salmon Commission 2024). Institutions responsible for publicly funded monitoring are ultimately accountable to ensure that resulting data are open and accessible in a timely manner.

Our analyses revealed numerous errors in NuSEDS that needed correction, including duplicate entries and mis-assigned population identifiers (undermining the efficacy of the slow-release approach discussed in the last paragraph). Data anomalies also included inconsistent use of zero and “NA” values, complicating efforts to discern whether monitoring occurred in these cases. Inconsistent use of zeroes and NAs hampers even simple investigations of monitoring efforts such as our analysis. Although many of these errors were easily corrected once identified, it is unclear if effort is made (or if sufficient capacity exists) to improve quality across the dataset. At a minimum, a transparent forum for documenting data errors (and ideally a mechanism to correct those errors) is needed.

As monitoring becomes increasingly distributed across DFO, First Nations, and non-governmental organisations (e.g., academic researchers, community groups, etc.), integrating data into NuSEDS will require increased investment in capacity to review and clean data to respect investment in monitoring (see Recommendation #2). Being more willing to share “preliminary” data, embracing crowd-source error checking, instituting a streamlined mechanism to correct identified errors, and implementing modern version control could help moderating the investment required.

### Recommendation #5: Work to develop data sharing protocols that facilitate data use while respecting Indigenous Data Sovereignty

FAIR data are Findable, Accessible, Interoperable, and Reusable - a set of principles that has been lauded by the scientific community for almost a decade (Wilkinson et al. 2016). More recently, the CARE principles for Indigenous Data Governance have also been developed to address concerns around Indigenous Data Sovereignty. CARE stands for data that are used for **collective** benefit, where Indigenous People maintain their **authority** to control their data, non-Indigenous institutions and individuals share a **responsibility** to engage respectfully, and data are applied following Indigenous Peoples’ **ethics** (Carroll et al. 2021). These two sets of principles can be complementary, and given that salmon are core to Indigenous livelihoods and cultures and salmon data are often collected by First Nations staff or members, both FAIR and CARE principles are applicable here (Carroll et al. 2021).

Facilitating access to salmon data requires understanding the hesitations and concerns of the data owners and custodians and this must be approached contextually. Fear of misuse or misinterpretation of data is an oft-cited explanation from DFO for limiting access to data, but ensuring that data are well documented with appropriate caveats and limitations is a logical antidote that is supported by existing government policy (Fisheries and Oceans Canada 2013). The Pacific Salmon Commission Fraser River Panel’s online tools for viewing and downloading both in-season and annual data on Fraser pink and sockeye abundance and timing (Pacific Salmon Commission 2024) demonstrate that timely, transparent, and appropriately caveated sharing of data is entirely feasible given appropriate resourcing. Rather than causing misunderstanding, publicly distributed data provide common baseline information that is foundational for equitable discussions about salmon management (Walters et al. 2008).

## Conclusions

Salmon are a foundational species in Pacific Canada, and counting them each year as they return home to spawn is integral to assessing and tracking biological status, managing local and regional fisheries, estimating the impact of myriad stressors, and evaluating recovery efforts. With the ongoing decline of monitoring salmon populations, both realised and potential knowledge gained from decades of counting salmon is eroding. Until now, conventional monitoring and management by DFO has focused on collecting the information needed to manage commercial harvest. By available public records, the proportion of salmon populations counted over the past decade (at the scale of individual populations spawning in their respective rivers/streams/lakes *or* the coarser scale of Conservation Units) is at the lowest it has been since the growth of counting programs in the 1950s. Pervasive salmon declines and the demise of the commercial salmon industry demand that we shift monitoring priorities to support the admirable blueprint set forth by the Wild Salmon Policy. As salmon distributions appear to be contracting, with pervasive declines at coarse scales (Connors et al. 2024), there is a need for *increased* monitoring of at-risk populations to avoid the undocumented erosion of salmon diversity. Revitalizing salmon spawner abundance monitoring - through redefined priorities, focused investments in salmon communities, and robust data practices - has the potential to advance shared goals and benefit both salmon themselves and the people and ecosystems that depend on them.

## Acknowledgements

We thank the Pacific Salmon Foundation for ongoing support of the Salmon Watersheds Program’s work to democratise salmon data, and for supporting staff to pursue this analysis. We thank Michael Price for reading an early draft of the manuscript and providing constructive feedback. Finally, we acknowledge and thank Dr. Jeff Hutchings, who has inspired our work, here and elsewhere, most clearly through his commitment to the integrity of the processes that connect science to decision makers in the context of fisheries.

## Author contributions

Conceptualization - EA, AB, EH, CA

Data curation - BC, SP

Formal analysis - BC, SP

Funding acquisition - KC, EA

Investigation - EA, BC, EH, AB, SP

Methodology - BC, EA, SP, EH, AB

Project administration – EA

Resources - EA, KC

Software - BC, SP, EA

Supervision - SP, KC

Validation - SP, BC

Visualization - BC, EA, SP

Writing - original draft - EA, SP, EH, CA, BC

Writing - review & editing - EA, SP, BC, EH, AB, KC

## Data availability statement

Cleaned spawner survey data are available at https://zenodo.org/records/14194638. Additional data and R code to reproduce all our analyses for this submission are archived at https://zenodo.org/records/14248905.

**Supplement A: Monitoring for fisheries or for fish? Declines in monitoring of salmon spawners continue despite a conservation crisis**

To access Supplement A, please follow this link: https://bookdown.org/salmonwatersheds/1_nuseds_collation_atkinson/1_nuseds_collation.html

**Supplement B: Monitoring for fisheries or for fish? Declines in monitoring of salmon spawners continue despite a conservation crisis**

To access Supplement B, please follow this link: https://bookdown.org/salmonwatersheds/atkinson_et_al_supp_main/

